# Alterations in auditory midbrain processing is observed in both female and male mouse model of Fragile X Syndrome

**DOI:** 10.1101/2025.06.04.657855

**Authors:** Abdullah Abdullah, Xiuping Liu, Kartikeya Murari, Jun Yan, Ning Cheng

## Abstract

**Introduction:** Sensory processing deficits presenting as auditory hypersensitivity is a common phenotype associated with Fragile X Syndrome (FXS), a leading monogenic cause of intellectual disability and autism spectrum disorder. Auditory hypersensitivity can also be observed in the *FMR1-*knockout (KO) mice, a well-established mouse model of autism and FXS. FXS is an X-linked disorder that is more prevalent in males compared to females, as a result most auditory and electrophysiology studies are performed in males. Previous *in-vivo* electrophysiology studies at the inferior colliculus (IC), an essential component of the auditory pathway, in male *FMR1-*KO mice at post-natal days 14, 21 and 34 (P14, P21 and P34) demonstrated increased neuronal firing, suggesting that the IC could play a role in auditory hypersensitivity. However, very little is known about the role of the central nucleus of the IC (ICc) and auditory hypersensitivity in females.

**Methods:** Here we investigated auditory processing at P20 and P30, representative of early and late developmental stage of auditory development, respectively, at the central nucleus of the IC (ICc) of both female and male *FMR1-*KO mice, compared with wildtype (WT) animals.

**Results:** *In-vivo* electrophysiology recordings from the ICc neurons of the *FMR1-*KO mice at both developmental ages demonstrated increased response magnitude measured by spike number to pure tones of varying frequency and amplitude, compared with age- and sex-matched WT animals. In addition, within the *FMR1-*KO group we also observed significant developmental and sex difference wherein higher response magnitudes were displayed at P20 and in the female mice. Minimum threshold of ICc neuron in the KO mice was significantly decreased at both P20 and P30. The ICc neurons in the KO mice also displayed increased response duration compared to WT animals at both P20 and P30, but significant sex difference was only observed at P30. Our data also indicated that the ICc neurons of both groups displayed weak negative relationship between latency and response magnitude at P20, and at P30 the WT mice showed a stronger relationship only in the female group.

In terms of developmental changes, we observed decreasing neuronal firing only in the KO mice between 20- and 30-day old female and male mice. Reduced response duration was observed in 30-day old mice of both sexes of both genotypes. Regarding minimum threshold, we observed a decline between early and late auditory development only in the male mice. Finally, our results also indicated that in the WT mice the reverse relationship between latency and response magnitude became more pronounced and consistent with age, a developmental trend that was absent in both the female and male *FMR1*-KO mice.

**Discussion:** Overall, our findings demonstrate that auditory processing deficits can also be observed in the female *FMR1*-KO mice using *in-vivo* electrophysiology studies, highlighting the importance of including female subjects in future studies. These results also indicate that auditory hypersensitivity can be observed robustly in younger mice, suggesting that the early development stage could be an ideal target for interventions.

## Introduction

Fragile X Syndrome (FXS) is a common monogenic cause of intellectual disability and autism spectrum disorder (ASD), with an estimated prevalence of 1 in 4000 males and 1 in 8000 females (Niu et al., 2017; Razak et al., 2021). The number of CGG trinucleotide repeats in the 5’ untranslated region of the Fragile X Messenger Ribonucleoprotein 1 (*FMR1*) gene varies within the general population, typically ranging between 6 to 55 repeats. When the CGG trinucleotide repeat number expands over 200 (a full mutation), hypermethylation occurs at the CpG site in the promoter region followed by the silencing of the *FMR1* gene (Liu et al., 2018; Zhou et al., 2016). Consequently, this disrupts the production of the Fragile X Messenger Ribonucleoprotein (FMRP) which leads to various symptoms in FXS like anxiety, social deficits, repetitive behavior and language impairments (Abbeduto et al., 2007; Niu et al., 2017).

Auditory hypersensitivity or intolerance to mild sounds is also a consistent symptom in individuals with FXS (Kokash et al., 2019; Tsiouris & Brown, 2004). In patients with FXS, event-related potential (ERP) manifests increased N1 and P2 amplitudes (Ethridge et al., 2019; Van Der Molen et al., 2012), indicating that auditory hypersensitivity can occur due to increased cortical responses to a sensory stimulus (Rotschafer & Razak, 2014). Moreover, auditory prepulse inhibition (PPI) of startles response, a behavioral study used to investigate sensorimotor gating (Graham, 1975), showed poor suppression of the auditory response to startle stimulus (Frankland et al., 2004; Hessl et al., 2009), indicating the presence of robust sensory processing deficits in FXS individuals (Sinclair et al., 2017).

The *FMR1-*knockout (KO) mouse is a well-established mouse model of FXS and syndromic autism (Kazdoba et al., 2014; Zhan et al., 2020), that share several FXS-phenotypes such as hyperactivity (Kazdoba et al., 2014) and alterations in dendritic spine shape and density (Grossman et al., 2006). The *FMR1-*KO mice also display noticeable sensory processing deficits like audiogenic seizures (AGS, typically occurring around postnatal day 20) (Gonzalez et al., 2019; Musumeci et al., 2000), altered sensorimotor gating (Chen & Toth, 2001; Kokash et al., 2019) and EEG/ERP abnormalities (Lovelace et al., 2018), modeling certain auditory phenotypes in humans with FXS. Physiological studies in the primary auditory cortex (AC) of *FMR1-*KO mice previously documented that auditory neurons were hypersensitive to acoustic stimuli when subjected to tones with varying frequencies and amplitudes (Wen et al., 2018).

The inferior colliculus (IC), also known as the auditory midbrain, is the convergence and integration center in the central auditory system (Liu, Wang, et al., 2024; Qi et al., 2020). Functionally, the inferior colliculus is anatomically subdivided into three major divisions namely the dorsal cortex (ICd), external cortex (ICe) and the central nucleus of the inferior colliculus (ICc) (Mansour et al., 2019). The ICc receives majority of the ascending auditory inputs through the lateral lemniscus (LL) which originates mainly from the cochlear nucleus (CN), lateral superior olive (LSO) and the nuclei of lateral lemniscus (LL) (Sitek et al., 2022). One study using post-natal day 18-24 (P18-24) mice observed that *FMR1* gene deletion in the VGLUT2-expressing glutamatergic neurons in subcortical brain regions such as the IC and thalamus was necessary and sufficient for the audiogenic seizure phenotype (Gonzalez et al., 2019). Another study observed elevated matrix metalloproteinase-9 (MMP-9) levels during early development (P7 and P12) at the IC along with prepulse inhibition deficits in young (P23-25) *FMR1-*KO mice (Kokash et al., 2019). Although these studies suggest that dysfunction of the IC neurons can play a role in auditory hypersensitivity, very little is known about whether the ICc neurons specifically is involved in this phenotype during early development.

FXS is an X-linked disorder (Razak et al., 2020) and is more prevalent in males than in females (Jacquemont et al., 2007). Although homozygous mutations are rare, female FXS individuals with heterozygous mutation can also present with severe symptoms like male FXS individuals possibly due to skewed X-chromosome inactivation (Heine-Suñer et al., 2003; Martorell et al., 2011). One study with both female and male FXS patients observed no significant difference in amplitude or latency in any of the event-related potential (ERP) components such as N1 or P2 (Knoth et al., 2014). However, another study with *FMR1-*KO mice observed enhanced ERP amplitudes in adult females compared to adult male *FMR1-*KO mice (Croom et al., 2024). Previously, a study by Nguyen et al demonstrated increased cFOS+ expression indicating elevated neuron activation at P21 and P34 and increased neuronal firing at the IC of P14, P21 and P34 *FMR1*-KO mice compared to controls (Nguyen et al., 2020). However, the study was performed only in males. Therefore, we hypothesized that the ICc neurons of both female and male *FMR1-*KO mice will display auditory hypersensitivity during development.

To test our hypothesis, we performed *in-vivo* electrophysiology recordings in response to pure tones of varying frequency and amplitude at the ICc of both female and male *FMR1-*KO mice during early (P20) and late (P30) development stages to quantify various response properties such as response magnitude, minimum threshold, relationship between latency and response magnitude, and response duration. Our results indicate that the ICc of the *FMR1*-KO mice display enhanced neuronal response **(Figure 1 E-H)**, prolonged response duration and decreased minimum threshold compared to the WT mice. The ICc neurons in both genotypes displayed weak inverse relationship between latency and response magnitude at P20, however at P30 the female WT mice displayed a stronger inverse relationship compared to the KO counterparts. We also observed significant sex differences within response magnitude in the *FMR1-*KO group and in response duration at P30.

**Figure 1:**
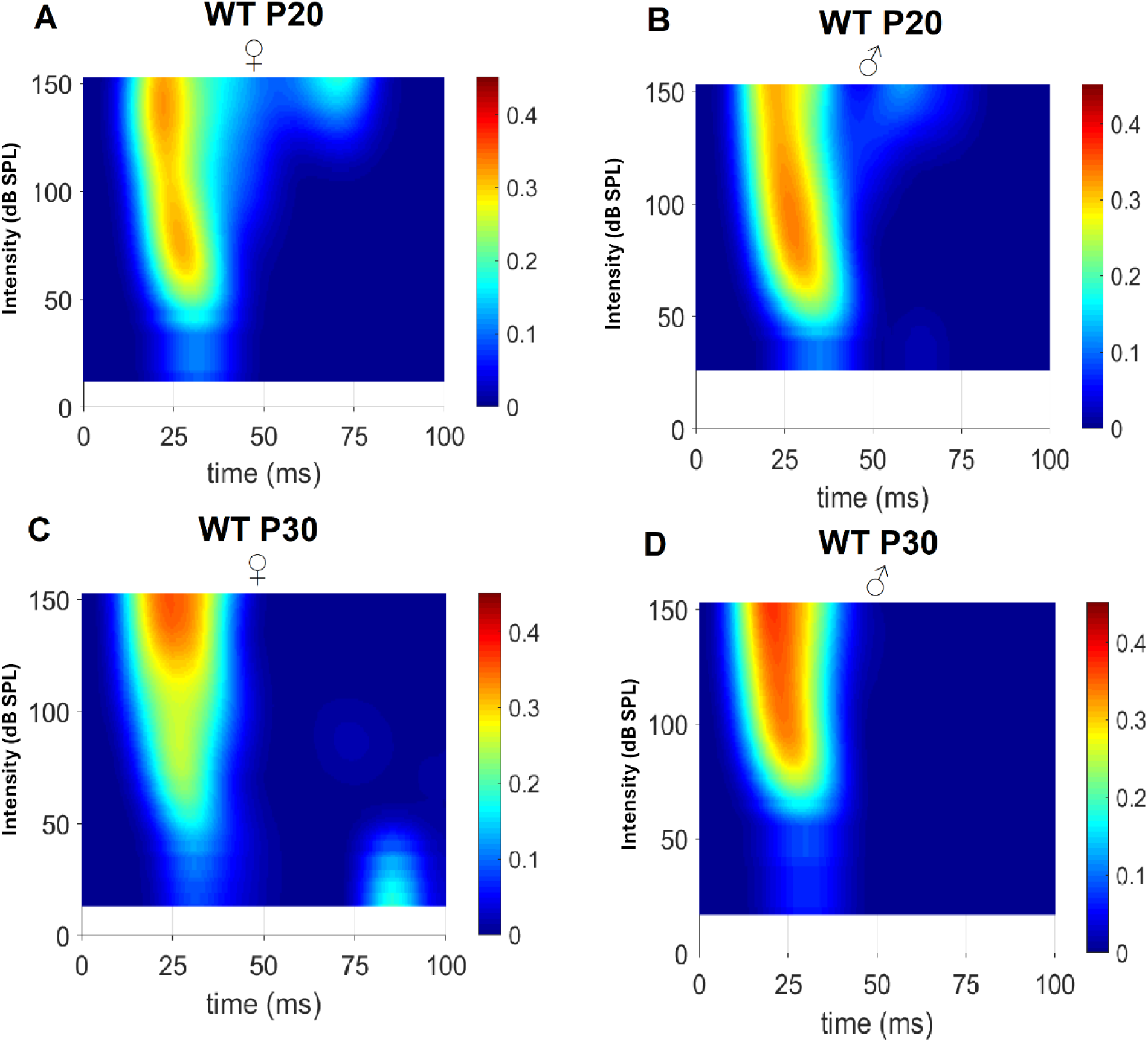
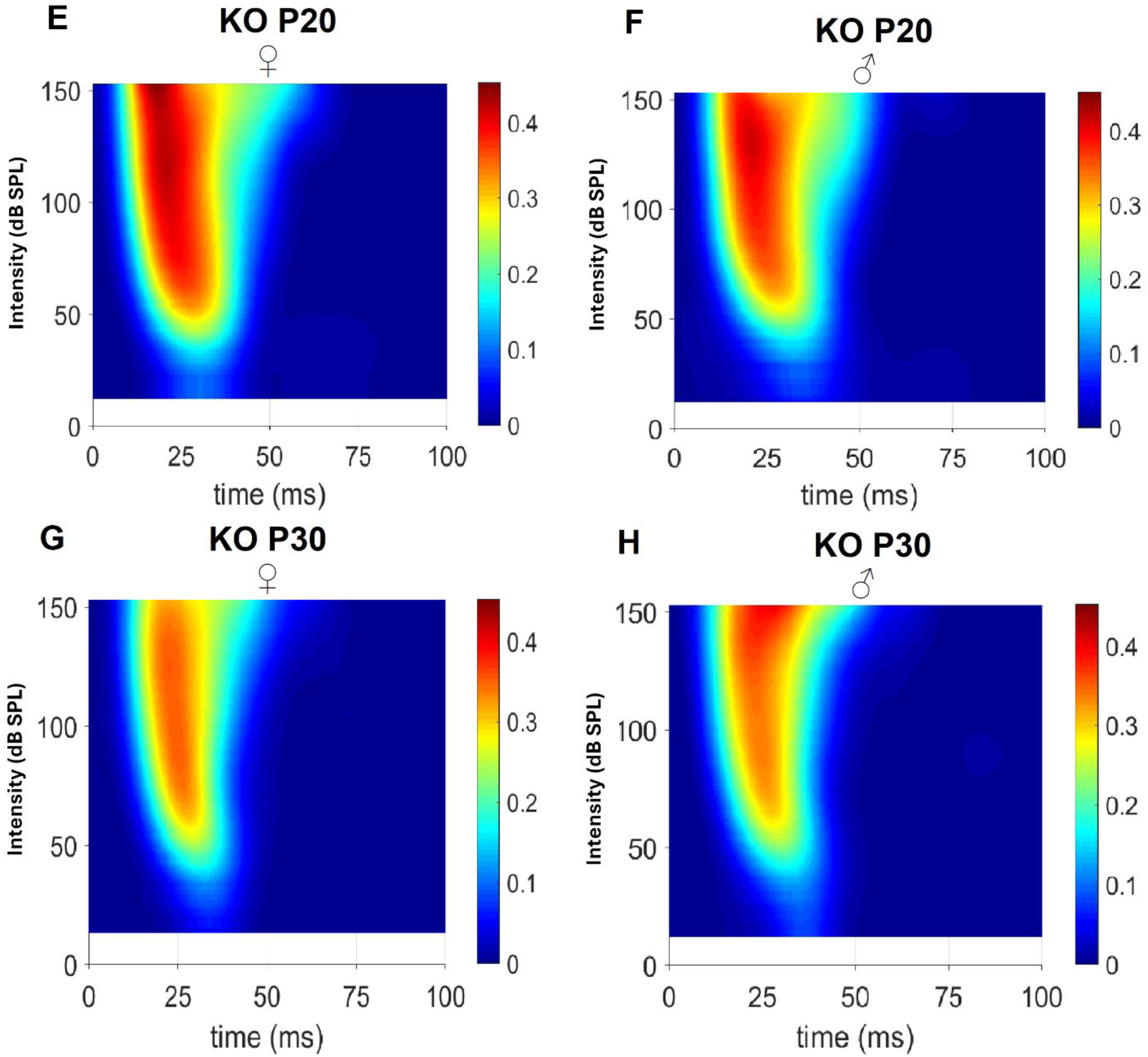
Contour plots of spike numbers with intensity at CF within 100ms recording duration. (A-D) Representative figures of a single WT unit at both sex and age. (E-H) Representative figures of a single KO unit at both sex and age. Color scale for each figure represents spike number at CF.

In terms of developmental changes, the response magnitudes of ICc neurons in both the female and male *FMR1-*KO mice decreased between P20 and P30. Response duration of both WT and KO female and male mice significantly diminished between early and late auditory development. Minimum threshold decreased with age only in the male mice. Finally, we observed that in the female and male WT group the inverse relationship between latency and response magnitude became more pronounced and consistent between 20- and 30-day old mice, a developmental trend that was not present in the KO group.

## Experimental Procedures

### Animals

Two mice colonies namely, wildtype (FVB.129P2-Pde6b+ Tyrc-ch/AntJ, Jax stock No: 004828) and *FMR1-*KO (FVB.129P2-*Pde6b*^+^ *Tyr^c-ch^ Fmr1^tm1Cgr^*/J, Jax stock No: 004624) were procured from Jackson Laboratory (ME, USA) and maintained in the mouse housing facility at the Health Sciences Animal Resources Centre, University of Calgary. Overall, a total of 202 units were recorded from the ICc of male postnatal day 20 (P20) WT (n= 21, from 5 mice), female P20 WT (n=23, from 4 mice), male P20 KO (n= 31, from 6 mice), female P20 KO (n= 14, from 4 mice), male P30 WT (n= 27, from 5 mice), female P30 WT (n= 19, from 4 mice), male P30 KO (n= 33, from 4 mice) and female P30 KO (n= 34, from 6 mice). The mice were bred in our animal facility at the University of Calgary. Mouse pups were weaned around post-natal day 20 and group-housed (up to five mice per cage) with same sex littermates. Mouse housing cages were kept at the facility with automated 12-hr light and dark cycle (lights on at 7:00 AM) with provisions to food (standard mouse chow) and water ad libitum. Electrophysiology recordings were performed between 09:00h and 19:00h. All procedures were carried out within the guidelines provided by the Canadian Council for Animal Care and was approved by the Health Sciences Animal Care Committee at the University of Calgary.

### Surgery

A mixture of ketamine (85 mg/kg, Bimeda-MTC Animal Health Inc., Canada) and xylazine (15 mg/kg, Bimeda-MTC Animal Health Inc., Canada) were injected intraperitoneally to anesthetize the mice throughout surgery and electrophysiological recording. Additional maintenance doses of ketamine and xylazine (17 and 3 mg/kg, respectively) were provided throughout this period, if the animal showed any responsiveness to tail pinching. Following anaesthesia, the mouse was placed on a feedback-controlled heating pad to maintain body temperature around 37°C throughout the surgery and recording period. The mouse’s head was fixed to a custom-made head holder by clamping between the palate and nasal bones, and the head-holder was adjusted until the Bregma-lambda of the skull aligned in the horizontal plane. The scalp, subcutaneous tissue and muscle were then carefully removed to expose the skull. A small craniotomy measuring 2 mm in diameter was performed using a dental drill to expose the inferior colliculus (IC, 0.5–2.0 mm posterior to the lambda, 0.5–2.0 mm left to the midline) based on coordinates from the Allen Brain Atlas. The electrophysiological recordings were all conducted in a sound-attenuated chamber with electromagnetic proofing.

### Recording in the ICc

Following surgery, a tungsten-wire microelectrode of ∼2-MΩ impedance was carefully lowered with a hydraulic microdrive into the left inferior colliculus. Action potentials received from the microelectrode were passed to a RA16PA multichannel preamplifier and further processed by a RA16 Medusa multichannel amplifier via an optic fiber cable (Tucker-Davis Technologies). Spike waveforms were saved using BrainWare acquisition software based on eight parameters including peak, valley, spike height, spike width, peak time, valley time and two voltage triggers. Tone-evoked responses from the ICc were generally observed when the microelectrode tip was located between 450 and 1000 μm in the inferior colliculus.

### Acoustic Stimulation

Tone bursts of 20 ms duration (5 ms rise and fall times) were digitally generated and converted to analog sinusoidal waves by an RZ6 MULTI I/O processor (Tucker-Davis Technologies, Inc., Gainesville, FL, USA). The analog signals were transferred through a digital attenuator and presented through a loudspeaker (MF1, Tucker-Davis Technologies., Gainesville, FL, USA) positioned 45° and 15 cm from the right ear of the mouse. Prior to the experiment, the tone amplitude (expressed as dB SPL, re. 20 μPa) of the loudspeaker was calibrated at the same position using a condenser microphone (Model 2520, Larson-Davis Laboratories, USA) and a microphone preamplifier (Model 2200C, Larson-Davis Laboratories, USA). BrainWare data acquisition software (Tucker-Davis Technologies, Inc., Gainesville, FL, USA) was used to manually or digitally deliver tones of different frequencies and intensities.

### Sampling receptive fields of ICc neurons

Once stable action potentials were observed, a frequency-amplitude scan (FA-scan) was performed to record the excitatory responses of the ICc neurons to tones of different frequency and intensity. The FA-scan consisted of a series of tones ranging from 14 to 74 dB SPL in 5 dB SPL steps, with frequencies equally spaced on a logarithmic scale. Each identical stimulus in the FA-scan was randomly delivered and repeated 5 times. The interstimulus interval between each tone bursts were 250 ms, and the entire session lasted for approximately 15 to 20 minutes. The receptive fields or frequency tuning curve for each ICc unit were obtained from the FA-scan data.

### Data acquisition and processing

The original action potential data acquired by BrainWare was archived in DAM files. These files were then processed by our custom-made data processing software, SoundCode. The trigger level was set manually at 20% greater than average background amplitude to detect tone-evoked neuronal firings or spikes, which could generally occur spontaneously, hence the trigger level allowed maximum exclusion of background noise in the signals. Neuronal firing pattern of multiple ICc units were plotted as raster, post-stimulus histogram (PST), and accumulated PST (PSTC) for quantification and further analysis. The following response properties were measured based on the tone evoked neuronal firing:

1. Spike number: Sum of spikes or action potential to identical stimulus within 0 to 100 ms from stimulus onset **(Figure 1)**.
2. Spike latency: The time interval from the onset of tone to the intersection point of the baseline and the rising slope of the PSTC.
3. Characteristic frequency (CF) and Minimum Threshold (MT): Both CF and MT were obtained based on the ICc receptive field. The CF is the single frequency at which the lowest response threshold (amplitude threshold) of the ICc unit is observed. The MT is intensity or amplitude at which the lowest response threshold of the neurons can be observed at characteristic frequency.
4. Response duration: Interval from the onset of stimulus to offset of neuronal response.

### Statistical Analysis

Statistical tests were performed in GraphPad Prism 10.2.3 (GraphPad Software, San Diego, California), and figures were generated in GraphPad Prism and MATLAB (MATLAB R2023b, MathWorks, Inc.). All statistical analyses are presented in the figure legends. All data is shown as Mean ± SEM and α was set at 0.05. For all response magnitude figures a two-way repeated measure ANOVA with post-hoc Tukey’s test, except for **Fig. 2I and J** where a three-way RM ANOVA, was performed. In addition, a simple linear regression analysis was performed to determine the relationship between latency and spike number. Finally, for all other analysis an ordinary two-way ANOVA test was performed wherever necessary, with subsequent post-hoc Tukey’s test.

**Figure 2:**
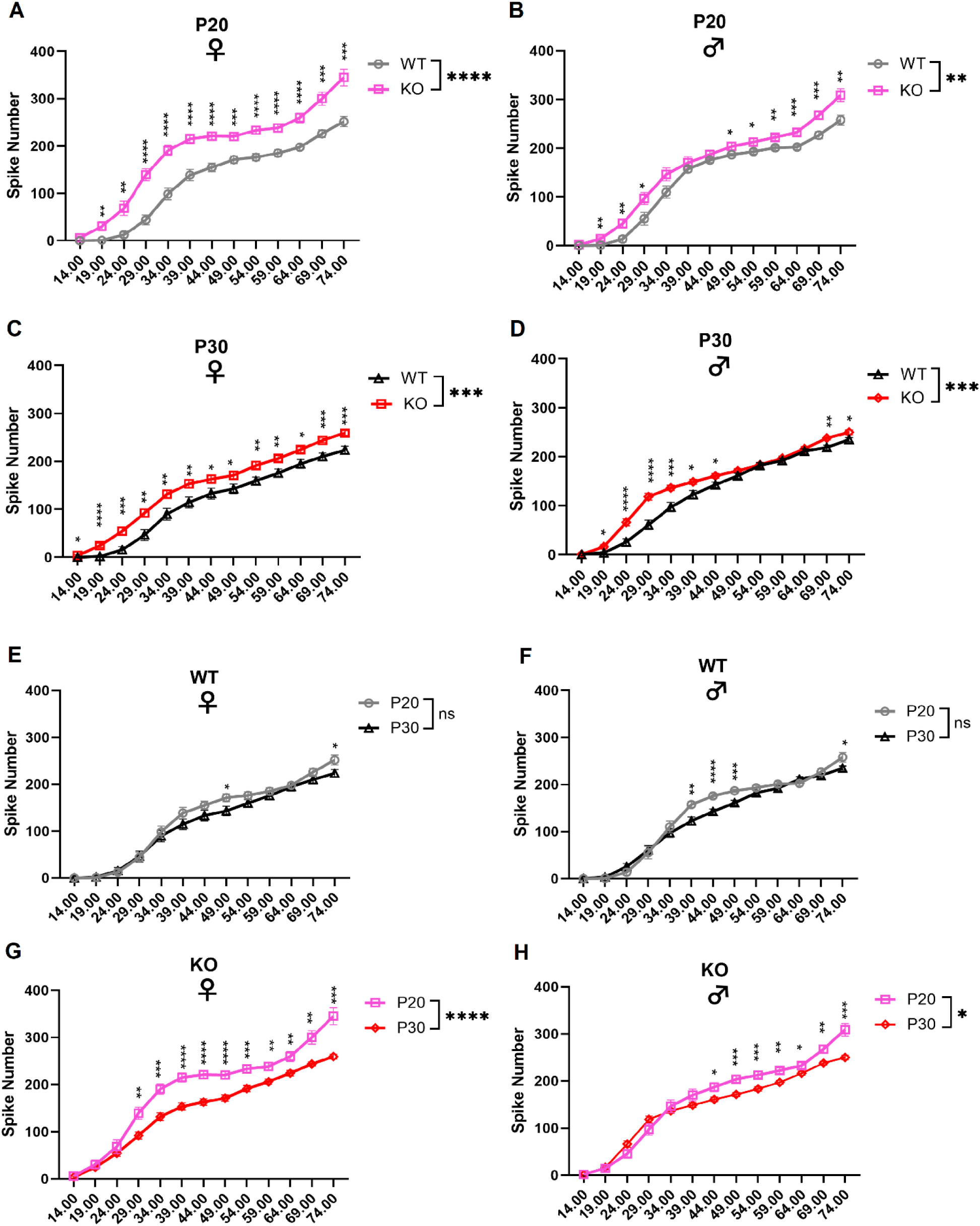
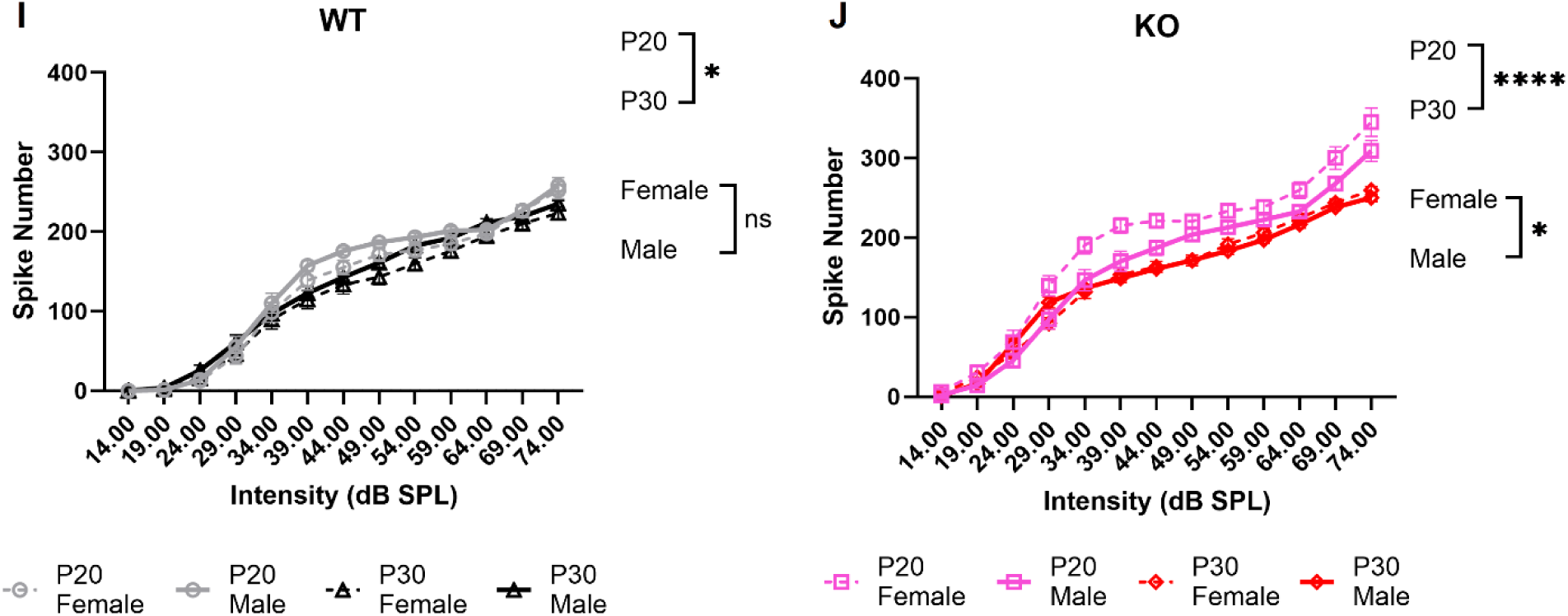
Response magnitude at different intensities in the ICc neurons of female and male *FMR1-*KO mice at CF. (A-D) Comparison between wild-type and *FMR1-*KO mice at both developmental timepoints and sex **(Table S1)**. (E-H) Comparison between P20 and P30 within genotype and sex. (I-J) Comparison of age and sex within the WT and KO groups respectively **(Table S2)**. Two-way RM ANOVA was performed for figures A-H, and three-way RM ANOVA was performed for figure I and J with post-hoc Tukey’s test (*:0.01, **: 0.001, ***:0.0001, ****: <0.0001).

## Results

### Neuronal response magnitude is enhanced at the ICc of female and male FMR1-KO mice

We first quantified response magnitude from *in-vivo* electrophysiology recordings at the ICc (**Figure 2**). At post-natal day 20 (P20), we observed that the response magnitude of *FMR1-*KO females was significantly higher (p= <0.0001) across all intensities compared to the wild-type (WT) mice. A similar trend was also observed at post-natal day 30 (P30), wherein significant genotype difference (p= 0.0002) was observed between female *FMR1-*KO and WT mice across all intensities. However, in males although there was significant genotype difference at P20 (p= 0.0034) and at P30 (p= 0.0009), the *FMR1-*KO mice did not exhibit increased response across all intensities **(Table S3)**.

To investigate the developmental changes, we also analyzed whether there were any age differences within genotype as shown in Figure 2 (**E-H**). Interestingly, we observed that there was significant age difference in both female (p= <0.0001) and male (p= 0.0308) *FMR1-*KO mice indicating that response magnitude was higher at P20 across most intensities **(Table S4)**. Finally, we also analyzed the overall sex and age difference within each genotype separately (**Figure 2I and J**). Response magnitude was higher in the female group compared to the male group in only the KO mice (p= 0.0126). Significant age difference was also observed within both the WT (p=0.0445) and the KO (p=<0.0001) mice. Overall, abnormal neuronal firing was enhanced in the *FMR1-*KO and especially in the female group during early development.

### Both female and male ICc neurons of FMR1-KO mice exhibited decreased minimum threshold

Hypersensitivity to a sensory stimulus is commonly observed in both female and male FXS patients (Lachiewicz et al., 2024). Therefore, we investigated whether ICc neurons of *FMR1-*KO mice can detect tones of lower intensity compared to WT mice by analyzing minimum threshold at best frequency (**Figure 3**). At both P20 and P30 (**Figure 3A and B**), there was significant genotype difference (p= <0.0001) but no sex difference. Post hoc pairwise comparison indicated that both female (P20, p= <0.0001; P30, p= <0.0001) and male (P20, p= 0.0012; P30, p= 0.0005) *FMR1-*KO displayed reduced minimum threshold compared to their respective WT counterparts. Finally, we found that only males showed a significant developmental difference (p= 0.0117) between P20 and P30, but not in the female group. Taken together, these data indicate that the ICc of both female and male *FMR1-*KO mice show increased sensitivity to tones of lower intensity, a response parameter that could develop differently in females and males.

**Figure 3:**
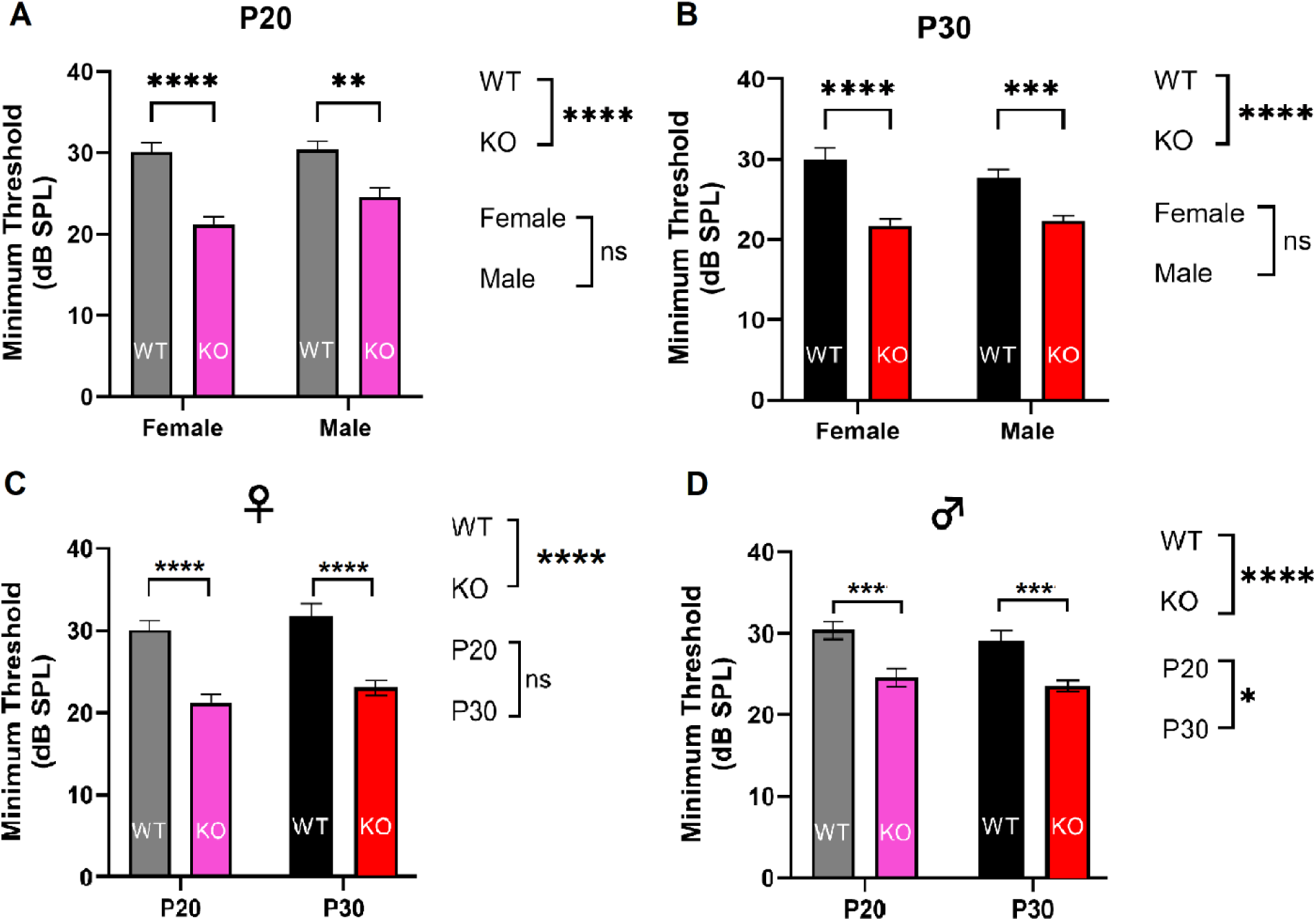
Minimum threshold of the ICc neurons of female and male *FMR1-*KO mice at CF. (A-B) Comparison between genotype and sex at both P20 and P30. (C-D) Difference between genotype and age in both the male and female groups. Ordinary two-way ANOVA was performed with post-hoc Tukey’s test for all above figures (*: 0.01, **: 0.001, ***:0.0001, ****: <0.0001)

### Weak inverse relationship between spike number and latency at the ICc of female and male P20 FMR1-KO mice

Based on our finding that neuronal response magnitude was enhanced in the *FMR1-*KO group, we also analyzed the relationship between latency and response magnitude to tones of 69.3 dB SPL, an intensity at which we previously observed robust responses irrespective of sex, age or genotype. Regarding differences between genotypes, significant difference between the slope (p= <0.0001) of regression lines was only observed in the P30 female mice (**Figure 4C**). The P30 female *FMR1-*KO mice displayed a shallower slope (b= -0.021) and poorer model fit (r^2^= 0.13) compared to the WT mice (b= -0.081, r^2^= 0.83) indicating a weaker relationship between latency and response magnitude in the KO mice. However, no other significant difference was observed in the other groups.

**Figure 4:**
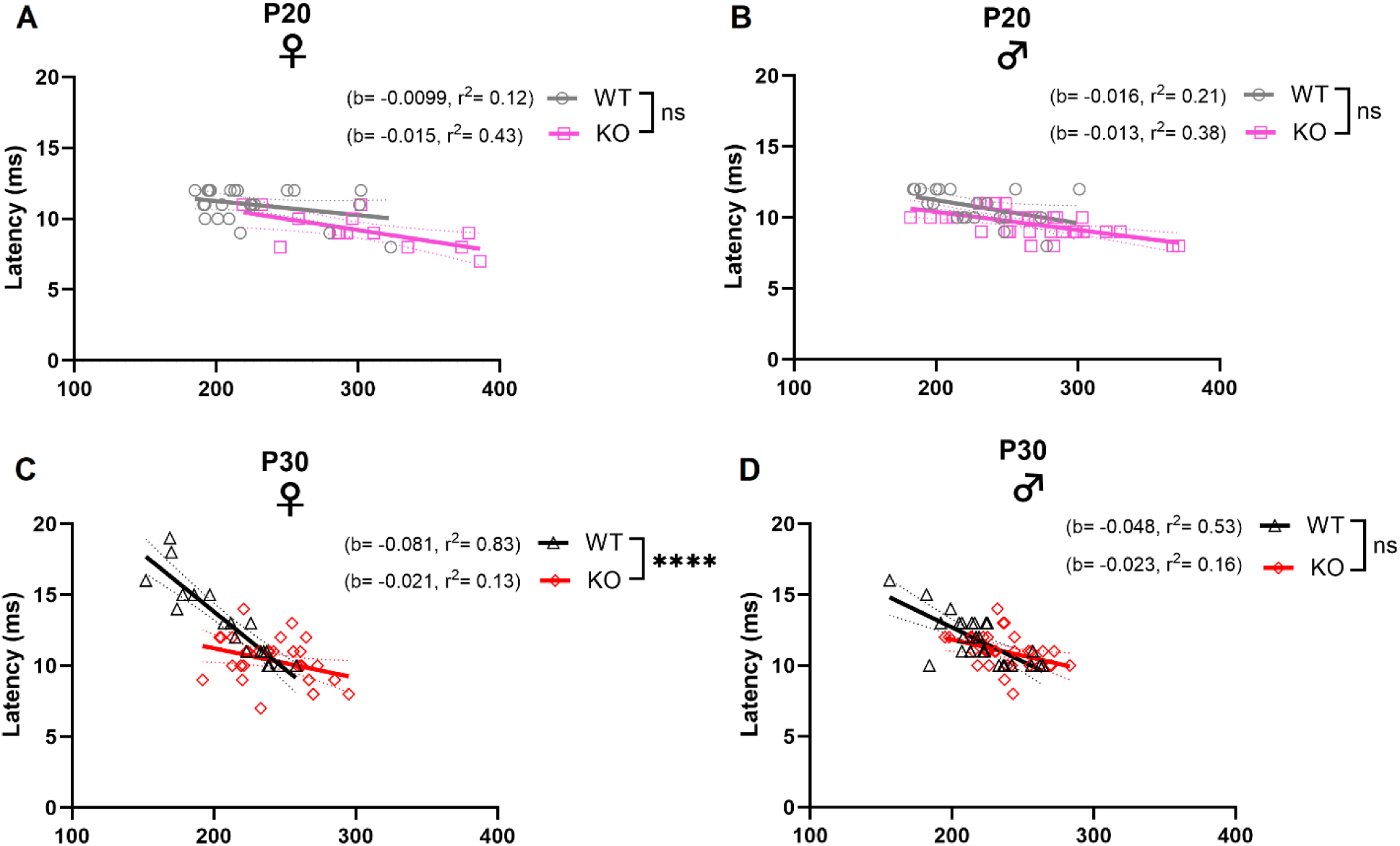
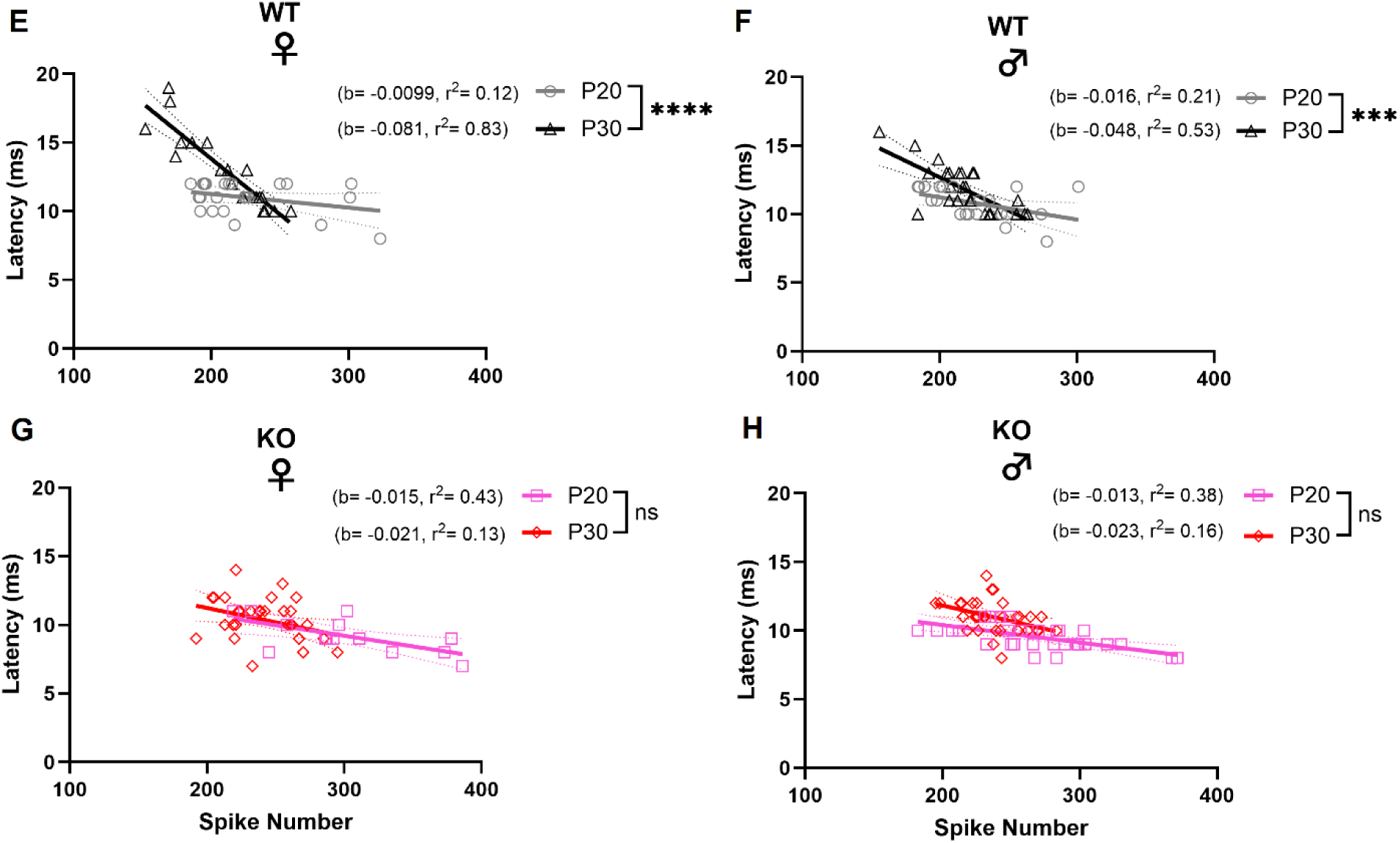
Relationship between latency (at CF and 69.3 dB SPL) and response magnitude at the ICc neurons of female and male *FMR1-*KO. (A-D) Comparison between WT and *FMR1-*KO at both developmental stages and sexes. (E-H) Comparison between P20 and P30 at in both genotypes and sexes. Line displayed with simple linear regression analysis.

In terms of developmental difference within genotype (**Figure 4E-H**), the regression lines were significantly different (p= <0.0001) in both female and male WT mice, which at P30 showed consistent (female, r^2^= 0.83; male, r^2^= 0.53) and pronounced negative relationship (female, b= -0.081; male, b= -0.048) compared to at P20 (female, b= -0.0099, r^2^= 0.12; male, b= -0.016, r^2^= 0.21), indicating that larger response magnitudes were more robustly associated with shorter latencies at the late development stage. However, no difference was observed between the regression lines in the KO group irrespective of sex, suggesting that the relationship between latency and response magnitude remains unchanged with age.

Collectively, our data indicates that auditory processing in the WT mice undergoes maturation indicated by the strengthening reverse relationship between latency and response magnitude, a developmental trend which is absent in the KO mice. As a result, all groups displayed weak negative relationships at P20, whereas at P30, the WT mice particularly the female group displayed stronger negative relation but not the KO mice.

### Longer response duration observed at the ICc of both female and male FMR1-KO mice

We also analyzed neuronal response duration (**Figure 5**) following tones of 69.3 dB SPL. Significant genotype difference (p= <0.0001) was observed at both P20 and P30, however sex difference was observed only at P30 (p= 0.0143). Furthermore, at both developmental timepoints neuronal response duration was significantly prolonged in both female (P20, p= 0.0082; P30, p= <0.0001) and male (P20, p= 0.0030; P30, p= 0.0138) *FMR1-*KO mice compared to their respective WT counterparts. Significant sex difference was only present at P30 (p= 0.0143), with KO female displaying increased response duration compared to the male mice (p= 0.0166).

**Figure 5:**
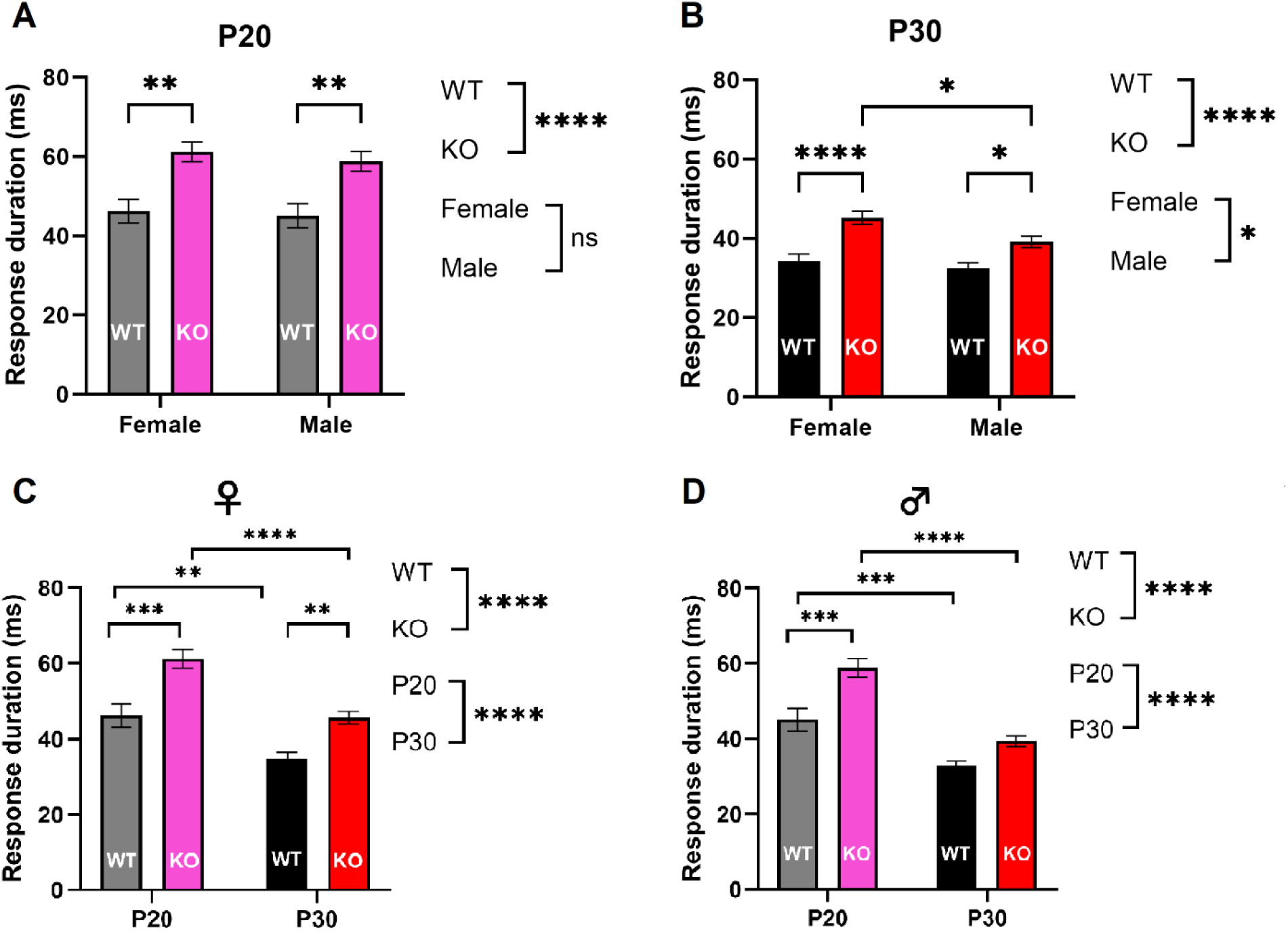
Response duration of ICc neurons of female and male *FMR1-*KO mice. (A-B) Comparison between genotype and sex at both P20 and P30. (C, D) Comparison between genotype and age in both the male and female groups. Ordinary two-way ANOVA with post-hoc Tukey’s test was performed for all above figures (*: 0.01, **: 0.001, ***:0.0001, ****: <0.0001).

Regarding developmental change **(Figure 5C and D)**, the response duration was significantly decreased (p= <0.0001) in both the female and male mice between P20 and P30. Notably, both the P30 WT (Female, p= 0.0031; Male, p= 0.0010) and *FMR1-*KO (Female, p= <0.0001; Male, p= <0.0001) mice exhibited decreased response duration compared to their P20 counterparts. As a result, our data suggests that both female and male *FMR1-*KO mice display increased response duration following stimulus onset. Moreover, response duration significantly decreased in both WT and KO mice with maturation.

## Discussion

We utilized *in-vivo* electrophysiology study and presented tones of different frequencies and amplitude to characterize the response properties of ICc neurons and established that increased sound sensitivity can be observed in both female and male *FMR1-*KO mice during development. One major finding at the ICc neurons was that the female FXS model also displayed auditory hypersensitivity, and the female *FMR1-*KO mice displayed increased neuronal firing compared to the male *FMR1-*KO mice. In addition, female *FMR1-*KO mice also displayed decreased minimum threshold, prolonged response duration and weaker negative relationship between latency and response magnitude compared to WT at P30.

We also found that, in general, atypical response properties were more evident at the ICc neuron of *FMR1-*KO mice at P20 compared to P30, suggesting that auditory hypersensitivity could develop at an early age which can persist to adulthood. Overall, our data suggests that the ICc is highly involved in auditory hypersensitivity in both the female and male *FMR1-*KO mice particularly during early development.

### Enhanced response magnitude in the *FMR1-*KO mice at both P20 and P30

Based on our *in-vivo* electrophysiology study, the ICc neurons of *FMR1-*KO mice consistently displayed increased response magnitudes especially in the young mice. Previous studies by Nguyen et al also observed elevated tone-evoked neuronal responses that declined with age at the ICc of only male *FMR1-*KO mice (Nguyen et al., 2020). However, our results indicate that the ICc neurons of both female and male *FMR1-*KO mice can display elevated neuronal responses to tones of different frequency and amplitude, compared to ICc neurons of age-matched wild-type mice. There was also a developmental age difference, with ICc neurons of *FMR1-*KO mice in P20 showing an increased response magnitude compared to P30 neurons, particularly in females within the *FMR1-*KO group.

One explanation for the increased sound sensitivity in the *FMR1-*KO mice could be attributed to the imbalance in the excitatory-inhibitory (E-I) network due to matrix metalloproteinase-9 (MMP-9), an FMRP regulated protein (Lovelace et al., 2020), that is elevated in the cortex and hippocampus in FXS individuals and the *FMR1-*KO mice (Christos et al., 2014; Lovelace et al., 2016). MMP-9 regulates the perineural net (PNN) surrounding parvalbumin-expressing (PV+) neurons in the IC, a population which includes both excitatory and inhibitory neurons. Notably, approximately one-quarter of PV+ neurons in the IC are GABAergic inhibitory neurons (Liu, Gao, et al., 2024), therefore MMP-9 plays a role in modulating cortical excitability (Razak et al., 2021).

One study observed increased MMP-9 levels at the IC only in young *FMR1-*KO mice (P7-P12), and reduction in MMP-9 levels reversed some of the sensory processing deficits (Kokash et al., 2019). Similarly, another study observed impaired development of PNN and PV along with elevated MMP-9 levels at the auditory cortex of P12-P18 male *FMR1-*KO mice. In addition, enhanced neuronal responses were observed using *in-vivo* electrophysiology recording which was reversed by reducing MMP-9 level (Wen et al., 2018). Taken together, we observed increased neuronal response at the central nucleus of the inferior colliculus of *FMR1-*KO mice prominently at P20, which suggests that increased MMP-9 levels could be involved in modulating neuronal hyper-responsiveness during early development.

### *FMR1-*KO mice show decreased minimum threshold at both P20 and P30

Nguyen et al observed that the minimum threshold of IC neurons in male *FMR1-*KO mice did not differ with the WT mice but declined significantly with age (Nguyen et al., 2020). Therefore, we quantified minimum threshold at characteristic frequency in female and male *FMR1-*KO mice at both developmental stages. Although we did not note any sex differences, there was significant genotype difference between the *FMR1-*KO and WT group at both P20 and P30. Both female and male *FMR1-*KO mice displayed decreased minimum threshold or sensitivity to tones of lower intensity compared to their WT counterparts. Another study by Ehret et al, investigated the development of response thresholds at the IC and observed a trend of decreasing threshold with age between post-natal days 10 and 20 (Ehret & Romand, 1992). Consistent with the previous study, we also observed significant age difference between P20 and P30 with decreasing minimum threshold, although only in the male group.

A behavioral study by Nielsen et al showed that the *FMR1-*KO mice consistently exhibited increased startle responses to lower intensity noise bursts (80 dB SPL) and decreased startle responses to higher intensity noise bursts (120 dB SPL) (Nielsen et al., 2002). Another study (Baker et al., 2010) which included both female and male *FMR1-*KO mice observed that auditory startle responses are lower at high tone intensities (105, 110 and 120 dB SPL). Our findings provide further evidence that dysfunction of the ICc neurons could play a role in the increased sensitivity and startle responses to tones of lower intensity in *FMR1-*KO mice.

### Latency against response magnitude relationship is weaker in the female *FMR1-*KO mice during late development

Another response parameter that we analyzed was the relationship between latency and response magnitude of the ICc neurons within genotype, age, and sex. We observed that particularly the P30 female *FMR1-*KO mice displayed weaker negative relationships compared to their WT counterparts, suggesting that the coupling between response magnitude and latency of response is disrupted in the KO mice. In general, neuronal responses occur at short latencies in the inferior colliculus (Syka et al., 2000) as observed in the P30 female WT mice, however impairments in neuronal processing could lead to variable response latencies in the KO mice. Although there are not many *in-vivo* electrophysiology studies that reported latency in FXS, auditory event-related potential (ERP) recordings from FXS individuals showed increased N2 component latencies (Van Der Molen et al., 2012). Similarly, ERP results from the auditory cortex (AC) in the FXS mouse model also observed prolonged N1 and P1 latencies (Lovelace et al., 2018).

The IC receives inhibitory inputs particularly by GABA_A_ receptors along with excitatory inputs which could play a role in regulating spike latency in the IC neurons (Wu et al., 2004). FMRP is highly expressed in GABA_A_ neurons (Gao et al., 2018), and loss of FMRP in FXS can lead to decreased expression of various GABA_A_ subunits and subsequent impairment of the GABA_A_ receptors (D’Hulst et al., 2006). One study (Gourévitch et al., 2020) observed an increase in peak latency of the response at the IC following blockade of GABA_A_ receptors suggesting that impaired GABA_A_ receptors activity could affect response latency in the IC.

In terms of developmental differences, we also observed that only the WT groups at P30 displayed stronger negative relationships, meaning that increased response magnitudes were observed more robustly at shorter latencies compared to the younger P20 mice. Age-related decrease in spike latency is consistent with reduction in the inhibitory drive to the IC (Simon et al., 2004), as demonstrated by a significant decrease in the number of GABA-energic neurons along with a low glutamate decarboxylase activity in the ICc with increasing age (Burianova et al., 2009; Ouda et al., 2015).

### Prolonged response duration at the ICc of *FMR1-*KO mice

We also studied the duration of the ICc neuron’s response which is the time between onset following the stimulus and the offset of the response. The ICc neurons of *FMR1-*KO mice exhibited significantly increased response durations at both P20 and P30 compared to the WT mice. There was also a sex difference with the female *FMR1-*KO mice displaying greater duration of response compared to the male *FMR1-*KO mice at P30. In terms of developmental change, both female and male *FMR1-*KO and WT mice showed decreasing response duration between P20 and P30. The FMRP protein regulates individual action potential of neurons via the BK channel during repetitive activity. In the absence of FMRP, BK channel activity reduces due to reduced interaction between FMRP and the β4 subunit of the BK channel leading to the broadening of individual action potential (Deng et al., 2013), possibly leading to sustained firing and prolonged neuronal response duration.

Alterations in inhibitory inputs due to GABA_A_ receptor activity deficits in FXS could also affect response duration, with one study showing that blocking GABA_A_ receptors could prolong response duration in ICc neurons (Gourévitch et al., 2020). Overall, our observation of prolonged response duration is a novel finding previously unreported in the *FMR1-*KO mice model and more electrophysiology studies are required to establish the mechanisms underlying prolonged response duration at the ICc.

### Sensory challenges in ASD and FXS

Individuals with ASD and FXS often experience a multitude of sensory challenges that can go beyond auditory hypersensitivity. Tactile hypersensitivity and atypical visual processing are common clinical features of ASD (Little, 2018; Puts et al., 2014), which are also observed in animal models of autism (Cheng et al., 2017; Cheng et al., 2020; Falcão et al., 2024). Similar sensory deficit phenotypes have also been reported in FXS patients (Cascio, 2010; Gallego et al., 2014) and in animal models (Arnett et al., 2014; Felgerolle et al., 2019). Therefore, these sensory processing deficits along with auditory hypersensitivity can pose significant social and behavioral challenges, underscoring the need for further research into the full spectrum of sensory difficulties to improve the quality of life of individuals with ASD and FXS.

## Conclusion

Auditory hypersensitivity is a prevalent phenotype in FXS. Previous studies in humans and rodent models observed abnormalities in cortical EEG/ERP recordings and *in-vivo* electrophysiology studies at the IC of *FMR1-*KO mice, but only in males. Our study highlights the importance of including females along with males in future FXS studies, since we demonstrated that sound sensitivity can be robustly observed at the ICc of females as in males, and differences between the sexes exist. Future research direction may involve carrying out further electrophysiology studies using different auditory paradigms to investigate the mechanisms that could potentially contribute to auditory hypersensitivity at the ICc of both female and male *FMR1-*KO mice. In addition, the developmental difference that we observed regarding response properties could also provide important information towards the development of early treatment guidelines unique to female and male FXS individuals. As a result, our findings not only are more inclusive and relevant to the wider FXS community but also indicates that early interventions could be beneficial to the quality of life of FXS individuals.

## Author Contribution

AA participated recording, data analysis and interpretation, writing and revising the manuscript. XL participated in designing the experiment, recording and data analysis. KM contributed to data analysis and interpretation. JY participated in designing the experiment, data analysis and interpretation, editing and revising the manuscript. NC designed the study, participated in data analysis and interpretation, and revised the manuscript.

## Supplementary material

aTables with statistical analysis data for corresponding figures provided as supplementary material.

## Abbreviations

FXS: Fragile X Syndrome
ASD: Autism spectrum disorder
ICc: Central nucleus of the inferior colliculus
*FMR1*: Fragile X Messenger Ribonucleoprotein 1
FMRP: Fragile X Messenger Ribonucleoprotein 1
*FMR1*-KO: *FMR1*-knockout, WT: Wildtype

