## Supplementary Table for "Alterations in auditory midbrain processing is observed in both female and male mouse model of Fragile X Syndrome"

### Supplementary material

### Tables

**Table S1:** Statistical analysis for **Figure 2 (A-D)** to compare response magnitude with different intensities at the ICc neurons of male and female wild-type and *FMR1*-KO mice at both developmental timepoints and sex. Statistical analysis: Two-way repeated measure ANOVA (p=*: 0.01, **: 0.001, ***:0.0001, ****: <0.0001, ns: non-significant).

| Figure | Source of Variation | P value | P value summary | F (DFn, DFd) |
|---|---|---|---|---|
| A | Genotype | <0.0001 | **** | F (1, 35) = 38.93 |
| | Intensity | <0.0001 | **** | F (4.010, 140.4) = 349.4 |
| | Genotype X Intensity | <0.0001 | **** | F (12, 420) = 6.448 |
| B | Genotype | 0.0034 | ** | F (1, 50) = 9.471 |
| | Intensity | <0.0001 | **** | F (3.254, 162.7) = 366.6 |
| | Genotype X Intensity | 0.0097 | ** | F (12, 600) = 2.222 |
| C | Genotype | 0.0002 | *** | F (1, 51) = 16.73 |
| | Intensity | <0.0001 | **** | F (4.177, 213.0) = 441.5 |
| | Genotype X Intensity | 0.0367 | * | F (12, 612) = 1.858 |
| D | Genotype | 0.0009 | *** | F (1, 58) = 12.20 |
| | Intensity | <0.0001 | **** | F (4.385, 254.3) = 800.3 |
| | Genotype X Intensity | <0.0001 | **** | F (12, 696) = 9.329 |

**Table S2:** Statistical analysis to compare response magnitude with different intensities at the ICc of male and female wild-type and *FMR1*-KO mice between P20 and P30 within genotype and sex **(Figure 2 E-H)**, and age and sex within each genotype **(Figure 2 I and J)**. Statistical analysis: Two- way repeated measure ANOVA was performed for **figures E-H** and three-way RM ANOVA was performed for **figures I and J** (p=*: 0.01, **: 0.001, ***:0.0001, ****: <0.0001, ns: non-significant).

| Group | Source of Variation | P value | P value summary | F (DFn, DFd) |
|---|---|---|---|---|
| E | Age | 0.2106 | ns | F (1, 40) = 1.619 |
| | Intensity | <0.0001 | **** | F (3.972, 158.9) = 365.7 |
| | Age X Intensity | 0.0428 | * | F (12, 480) = 1.818 |
| F | Age | 0.0924 | ns | F (1, 46) = 2.955 |
| | Intensity | <0.0001 | **** | F (3.324, 152.9) = 504.1 |
| | Age X Intensity | <0.0001 | **** | F (12, 552) = 4.101 |
| G | Age | <0.0001 | **** | F (1, 46) = 24.51 |
| | Intensity | <0.0001 | **** | F (4.047, 186.2) = 428.1 |
| | Age X Intensity | <0.0001 | **** | F (12, 552) = 7.303 |
| H | Age | 0.0308 | * | F (1, 62) = 4.886 |
| | Intensity | <0.0001 | **** | F (4.002, 248.1) = 549.5 |
| | Age X Intensity | <0.0001 | **** | F (12, 744) = 9.069 |
| I | Age | 0.0445 | * | F (1, 86) = 4.160 |
| | Sex | 0.0509 | ns | F (1, 86) = 3.919 |
| | Age X Sex X Intensity | 0.9737 | ns | F (12, 1032) = 0.3703 |
| J | Age | <0.0001 | **** | F (1, 108) = 26.41 |
| | Sex | 0.0126 | * | F (1, 108) = 6.436 |
| | Age X Sex X Intensity | 0.0023 | ** | F (12, 1296) = 2.567 |

**Table S3:** Post-hoc comparison for **Figure 2 (A-D)** to compare response magnitude with different intensities at the ICc of male and female wild-type and *FMR1*-KO mice at both developmental timepoints and sex. Statistical Analysis: Tukey's multiple comparison test (p= *: 0.01, **: 0.001, ***:0.0001, ****: <0.0001, ns: non-significant).

| Group | Intensity (dB SPL) | Mean Diff. | 95.00% CI of diff. | P Value Summary | Adjusted P Value |
|---|---|---|---|---|---|
| P20 Female | 14.85 | -6.143 | -15.16 to 2.876 | ns | 0.1649 |
| | 19.8 | -29.27 | -49.81 to -8.723 | ** | 0.0087 |

| | | | | | |
|---|---|---|---|---|---|
| WT vs KO | 24.75 | -55.75 | -89.11 to -22.39 | ** | 0.0026 |
| | 29.7 | -95.37 | -128.7 to -62.06 | **** | <0.0001 |
| | 34.65 | -91.66 | -125.5 to -57.79 | **** | <0.0001 |
| | 39.6 | -76.30 | -107.7 to -44.86 | **** | <0.0001 |
| | 44.55 | -66.01 | -91.70 to -40.33 | **** | <0.0001 |
| | 49.5 | -49.01 | -72.23 to -25.79 | *** | 0.0001 |
| | 54.45 | -56.70 | -80.59 to -32.82 | **** | <0.0001 |
| | 59.4 | -53.44 | -75.99 to -30.89 | **** | <0.0001 |
| | 64.35 | -61.93 | -87.24 to -36.63 | **** | <0.0001 |
| | 69.3 | -73.87 | -108.0 to -39.75 | *** | 0.0002 |
| | 74.25 | -93.27 | -136.4 to -50.13 | *** | 0.0002 |
| P20 Male WT vs KO | 14.85 | -1.710 | -4.160 to 0.7403 | ns | 0.1644 |
| | 19.8 | -13.42 | -21.75 to -5.088 | ** | 0.0025 |
| | 24.75 | -31.63 | -53.81 to -9.451 | ** | 0.0061 |
| | 29.7 | -41.59 | -77.58 to -5.597 | * | 0.0245 |
| | 34.65 | -36.28 | -73.09 to 0.5325 | ns | 0.0533 |
| | 39.6 | -12.72 | -40.58 to 15.15 | ns | 0.3623 |
| | 44.55 | -11.30 | -32.19 to 9.584 | ns | 0.2811 |
| | 49.5 | -17.11 | -31.94 to -2.275 | * | 0.0249 |
| | 54.45 | -19.81 | -36.56 to -3.055 | * | 0.0214 |
| | 59.4 | -21.52 | -37.20 to -5.832 | ** | 0.0082 |
| | 64.35 | -30.63 | -46.41 to -14.86 | *** | 0.0003 |
| | 69.3 | -41.24 | -62.75 to -19.72 | *** | 0.0003 |
| | 74.25 | -51.06 | -84.14 to -17.99 | ** | 0.0032 |
| P30 Female WT vs KO | 14.85 | -3.824 | -7.252 to -0.3947 | * | 0.0300 |
| | 19.8 | -22.38 | -32.27 to -12.50 | **** | <0.0001 |
| | 24.75 | -38.83 | -58.21 to -19.44 | *** | 0.0002 |
| | 29.7 | -45.81 | -73.62 to -18.00 | ** | 0.0020 |
| | 34.65 | -41.95 | -72.34 to -11.56 | ** | 0.0083 |
| | 39.6 | -38.85 | -67.07 to -10.63 | ** | 0.0084 |

| | | | | | |
|---|---|---|---|---|---|
| | 44.55 | -29.84 | -57.20 to -2.471 | * | 0.0336 |
| | 49.5 | -28.17 | -52.65 to -3.677 | * | 0.0254 |
| | 54.45 | -31.96 | -52.93 to -10.99 | ** | 0.0037 |
| | 59.4 | -30.42 | -50.99 to -9.841 | ** | 0.0050 |
| | 64.35 | -29.62 | -52.21 to -7.031 | * | 0.0118 |
| | 69.3 | -33.62 | -50.77 to -16.47 | *** | 0.0004 |
| | 74.25 | -35.93 | -55.44 to -16.43 | *** | 0.0007 |
| P30 Male WT vs KO | 14.85 | -0.3098 | -2.132 to 1.512 | ns | 0.7348 |
| | 19.8 | -12.68 | -22.26 to -3.097 | * | 0.0106 |
| | 24.75 | -40.29 | -59.56 to -21.03 | **** | <0.0001 |
| | 29.7 | -57.64 | -81.66 to -33.62 | **** | <0.0001 |
| | 34.65 | -39.26 | -61.03 to -17.48 | *** | 0.0008 |
| | 39.6 | -26.03 | -46.14 to -5.918 | * | 0.0124 |
| | 44.55 | -18.05 | -32.59 to -3.515 | * | 0.0159 |
| | 49.5 | -10.15 | -23.89 to 3.593 | ns | 0.1445 |
| | 54.45 | -0.8687 | -14.94 to 13.20 | ns | 0.9020 |
| | 59.4 | -5.051 | -18.78 to 8.681 | ns | 0.4644 |
| | 64.35 | -4.838 | -18.32 to 8.642 | ns | 0.4752 |
| | 69.3 | -19.06 | -31.32 to -6.806 | ** | 0.0030 |
| | 74.25 | -14.99 | -27.84 to -2.129 | * | 0.0231 |

**Table S4:** Post-hoc comparison for **Figure 2 (E-H)** to compare response magnitude with different intensities at the ICc of male and female wild-type and *FMR1*-KO mice between P20 and P30 within genotype and sex. Statistical Analysis: Tukey's multiple comparison test (p= *: 0.01, **: 0.001, ***:0.0001, ****: <0.0001, ns: non-significant).

| Group | Intensity (dB SPL) | Mean Diff. | 95.00% CI of diff. | P Value Summary | Adjusted P Value |
|---|---|---|---|---|---|
| WT Female | 14.85 | 0.000 | | | |
| | 19.8 | -0.9588 | -6.299 to 4.381 | ns | 0.7162 |

| | | | | | |
|---|---|---|---|---|---|
| P20 vs P30 | 24.75 | -2.963 | -20.56 to 14.63 | ns | 0.7347 |
| | 29.7 | -2.291 | -33.74 to 29.16 | ns | 0.8836 |
| | 34.65 | 8.854 | -26.98 to 44.69 | ns | 0.6202 |
| | 39.6 | 24.18 | -9.607 to 57.96 | ns | 0.1559 |
| | 44.55 | 21.76 | -7.869 to 51.39 | ns | 0.1450 |
| | 49.5 | 28.19 | 2.412 to 53.97 | * | 0.0329 |
| | 54.45 | 16.97 | -5.218 to 39.15 | ns | 0.1300 |
| | 59.4 | 9.341 | -13.78 to 32.47 | ns | 0.4188 |
| | 64.35 | 2.993 | -19.84 to 25.82 | ns | 0.7911 |
| | 69.3 | 15.87 | -6.297 to 38.03 | ns | 0.1557 |
| | 74.25 | 28.45 | 1.390 to 55.51 | * | 0.0398 |
| WT Male P20 vs P30 | 14.85 | -0.6296 | -1.924 to 0.6646 | ns | 0.3265 |
| | 19.8 | -2.926 | -7.903 to 2.051 | ns | 0.2408 |
| | 24.75 | -11.94 | -30.51 to 6.638 | ns | 0.2022 |
| | 29.7 | -5.339 | -38.23 to 27.56 | ns | 0.7444 |
| | 34.65 | 12.98 | -18.59 to 44.55 | ns | 0.4108 |
| | 39.6 | 34.72 | 13.80 to 55.64 | ** | 0.0017 |
| | 44.55 | 32.87 | 18.65 to 47.09 | **** | <0.0001 |
| | 49.5 | 25.48 | 13.27 to 37.70 | *** | 0.0001 |
| | 54.45 | 10.54 | -4.237 to 25.32 | ns | 0.1578 |
| | 59.4 | 8.778 | -5.258 to 22.81 | ns | 0.2144 |
| | 64.35 | -9.222 | -23.14 to 4.691 | ns | 0.1886 |
| | 69.3 | 7.667 | -9.805 to 25.14 | ns | 0.3796 |
| | 74.25 | 22.93 | 0.6595 to 45.19 | * | 0.0440 |
| KO Female P20 vs P30 | 14.85 | 2.319 | -7.162 to 11.80 | ns | 0.6129 |
| | 19.8 | 5.924 | -15.89 to 27.74 | ns | 0.5761 |
| | 24.75 | 13.95 | -20.24 to 48.14 | ns | 0.4034 |
| | 29.7 | 47.26 | 17.21 to 77.32 | ** | 0.0036 |
| | 34.65 | 58.56 | 30.55 to 86.58 | *** | 0.0002 |
| | 39.6 | 61.63 | 36.34 to 86.91 | **** | <0.0001 |

| | | | | | |
|---|---|---|---|---|---|
| | 44.55 | 57.94 | 34.99 to 80.89 | **** | <0.0001 |
| | 49.5 | 49.03 | 27.29 to 70.78 | **** | <0.0001 |
| | 54.45 | 41.71 | 18.91 to 64.51 | *** | 0.0008 |
| | 59.4 | 32.37 | 12.41 to 52.32 | ** | 0.0025 |
| | 64.35 | 35.30 | 10.20 to 60.41 | ** | 0.0081 |
| | 69.3 | 56.12 | 24.52 to 87.71 | ** | 0.0017 |
| | 74.25 | 85.79 | 46.18 to 125.4 | *** | 0.0003 |
| KO Male P20 vs P30 | 14.85 | 0.7703 | -1.982 to 3.522 | ns | 0.5760 |
| | 19.8 | -2.187 | -13.76 to 9.382 | ns | 0.7068 |
| | 24.75 | -20.60 | -43.37 to 2.169 | ns | 0.0753 |
| | 29.7 | -21.39 | -49.71 to 6.928 | ns | 0.1354 |
| | 34.65 | 10.00 | -19.18 to 39.19 | ns | 0.4923 |
| | 39.6 | 21.41 | -5.884 to 48.71 | ns | 0.1208 |
| | 44.55 | 26.13 | 5.013 to 47.24 | * | 0.0164 |
| | 49.5 | 32.44 | 16.35 to 48.54 | *** | 0.0002 |
| | 54.45 | 29.48 | 13.32 to 45.64 | *** | 0.0006 |
| | 59.4 | 25.24 | 9.814 to 40.67 | ** | 0.0018 |
| | 64.35 | 16.57 | 1.160 to 31.99 | * | 0.0356 |
| | 69.3 | 29.84 | 12.07 to 47.62 | ** | 0.0015 |
| | 74.25 | 59.00 | 30.76 to 87.24 | *** | 0.0001 |
